# Decoding substrate specificity determining factors in glycosyltransferase-B enzymes – Insights from machine learning models

**DOI:** 10.1101/2024.11.26.625515

**Authors:** Samantha G. Hennen, Yannick J. Bomble, Breanna R. Urbanowicz, Vivek S. Bharadwaj

**Author notes:** **Corresponding Author** Vivek Bharadwaj.

## Abstract

Substrate specificity is an essential characteristic of any enzyme’s function and an understanding of the factors that determine this specificity is crucial for enzyme engineering. Unlike the structure of an enzyme which is directly impacted by its sequence, substrate specificity as an enzyme attribute involves a rather indirect relationship with sequence as it also depends on structural aspects that dictate substrate accessibility and active site dynamics. In this study, we explore the performance of classifier-based machine learning models trained on curated sequence and structural data for a class of glycosyltransferases (GTs), namely GT-Bs, to understand their substrate specificity determining factors. GTs enable the transfer of sugar moieties to other biomolecules such as oligosaccharides or proteins and are found in all kingdoms of life. In plants, GTs participate in the biosynthesis of plant cell wall biopolymers (eg: hemicelluloses and pectins) and are an integral part of the enzymatic machinery that enables the storage of carbon and energy as plant biomass. To elucidate the substrate specificity of uncharacterized GT-Bs, we constructed multi-label machine learning models (Support Vector Classifier, K-Nearest Neighbors, Gaussian Naïve-Bayes, Random Forest) that incorporate both sequence and structural features. These models achieve good predictive accuracies on test datasets. However, despite our use of structural information, we highlight that there is further scope for improvement in training these models to draw interpretable relationships between sequence, structure and substrate specificity determining motifs in GT-Bs.

## 1. Introduction

Plants employ amongst Nature’s most efficient carbon capture mechanisms, storing much of the world’s carbon, and are therefore a valuable resource for conversion to fuels and products. The polysaccharides present in the cell walls that make up the bulk of plant biomass are synthesized, constructed, and modified by a conglomerate of enzymes. A thorough understanding of these enzymes, their substrate specificity and catalytic functions is of utmost importance for our fundamental knowledge of how the carbon fixed via photosynthesis is converted and stored in plant biomass, and for facilitating the technology-development to design tailored biopolymers for materials.^1–4^ Among the various classes of enzymes involved in plant cell wall biosynthesis, of particular interest are glycosyltransferases (GTs) that catalyze the formation of glycosidic bonds by transferring sugar moieties from sugar-nucleotide donors to oligosaccharide acceptors.^5, 6^ These enzymes are responsible for the formation of complex glycopolymers that constitute a large portion of the cell wall governing its architecture.^5^

GTs are ubiquitous in both plant and animal species and have thus far been classified into 117 families in the CAZy database on the basis of their sequence similarity.^5,7^ GTs are known to adopt one of three major structural folds: GT-A, -B, and -C and comprise 21, 27, and 10 families identified in the CAZy database respectively.^7^ Unfortunately, the structural folds of several families have not yet been officially classified by the CAZy database^7^, due to the lack of experimental evidence. While individual members of some families have been proposed to adopt certain folds, such as the fucosyltransferase *At*FUT1 from the GT37 family,^8^ the structural, functional and mechanistic details of the majority of GTs are still not well understood. The GT-B fold was first identified as a distinct folding superfamily in 2001, and is characterized by a catalytic site localized between two Rossmann-like subdomains.^9^ GT-Bs are of particular interest for their important role in the synthesis of non-cellulosic plant polysaccharides, a major portion of plant biomass.^5^ GT-B enzymes in the GT37 and GT47 families, for example, are key enzymes involved in xyloglucan synthesis, the major hemicellulose in the primary cell walls of dicots.^8, 10, 11, 12^ Despite their important role in plants and the presence of a diverse range of monosaccharides in plants, detailed knowledge of the substrate specificities of many GT-B proteins is yet to be fully understood.

Current approaches used to investigate the molecular mechanisms of biocatalysts often rely on the use of experimentally determined protein crystal structures to develop functional hypotheses that form the basis for subsequent biochemical and mechanistic investigations. While these structures offer highly detailed information, they can be difficult and tedious to generate even for a single candidate.^13, 14–17^ This approach becomes intractable for a comprehensive exploration of the impact of mutations or natural variants on substrate specificities in an enzyme family or class. Fortunately, thanks to advances in genomics, there is currently a wealth of available sequence data, allowing for machine learning (ML) approaches that can find patterns throughout large numbers of sequences and relate it to substrate specificity.^18,19, 20^ Additionally, there are now over 200 million predicted protein structures available in the AlphaFold2 (AF2) database, allowing for structural information for all protein coding genes from a multitude of species to be easily considered alongside sequence data.^21,22^

Machine learning approaches that leverage this abundance of data are being increasingly used to elucidate properties and activities of enzymes and have been employed in recent studies to predict substrate specificity for some GTs. Yang et al. generated an activity assay for GT-1 enzymes, which is a family known to adopt a GT-B fold form, and used decision trees trained on this data to predict donor and acceptor substrate specificity.^23^ Taujale et al. evaluated the use of tree-based models to predict the donor specificities of GT-A sequences, the most abundant and well-characterized of the three GT folds.^24^ There has been no significant efforts focused on predicting the specificities across diverse GT families that adopt a GT-B fold. Predicting substrate specificity is a challenging task. Amongst the multitude of factors that determine substrate specificity, some are directly related to sequence eg: structure, while others are consequent attributes of sequence eg: active site dynamics, and many more are totally sequence-independent eg: substrate chemical environments.^25^ Furthermore, many GT families display polyspecificity, with GT47 being a key example wherein members utilize a variety of donors and acceptors, making precise functional predictions even more difficult and unreliable. It is especially challenging to connect sequence to specificity for GT-Bs, as this fold family has little inter-family sequence similarity, and multiple families lack experimentally characterized structures.^24,26^

In this study, we curate sequence-activity data on GT-Bs, build classifier ML models, compare the performance of four types of models (Random Forest, Support Vector Machines, and K-Nearest Neighbors) on their ability to predict their donor binding specificity, and attempt to connect critical residue features identified from the ML models to the enzyme structure to identify substrate specificity determining motifs in GT-Bs. Our approach began with curating GT-B sequences for training and testing our models and consisted of sequences obtained from the CaZY and Uniprot databases that have been annotated for activity on one of seven diphosphate sugar substrates (GDP-Mannose, GDP-Fucose, UDP-Galactose, UDP-Glucose, UDP-Glucuronic acid, UDP-Xylose, UDP-Rhamnose). This was followed by the description of these GT-B sequences in terms of sequence features such as residue-based polarity, hydrophobicity, and charge, as well as structural features, including solvent accessible surface area and secondary structure (obtained from AF2 predicted structures) to build our ML models. Additionally, due to the large diversity within GT-B families in sequence and structure, only the residues in the Rossmann-like subdomains and catalytic site in the AF2 structural models were aligned and featurized. This was done to generate more meaningful multiple sequence alignments (MSAs), as well as focus the model on positions more likely to dictate substrate specificity. Furthermore, unlike previous studies, these models have been built to predict multiple substrates for each enzyme, as this is an important consideration for GT families that are known to utilize multiple donor substrates, such as GT1, GT4, GT31, and GT47. The trained models have proved to be accurate with the KNN model achieving cross-validation and test scores of 95% and 85%, respectively. We then perform a conservation analysis on the set of residues whose features contribute most to decision-making in the model and translate it to the enzyme structure and results of docking calculations to gauge their importance in substrate-binding.

## 2. Methods

### 2.1. Dataset collection

The training dataset comprises GT-B enzymes from 135 species and subspecies gathered from the UniProt database (SI Figure 1). To confirm GT-B identity, only families previously shown to adopt a GT-B fold, as indicated in the Carbohydrate Active Enzymes (CAZy) database, were included in the dataset. Further, only UniProt-reviewed proteins were considered to ensure high quality data.^7^ UniProt catalytic activity information was mined to determine donor substrate specifity, with proteins utilizing a donor substrate known to participate in plant carbohydrate biosynthesis selected for the dataset. Five GT47, four GT37, and five GT61 enzymes were also added, as they are known to adopt a GT-B fold, with their substrates were identified from previous literature and the CAZy database.^7, 8, 10, 11, 13, 27, 28, 29, 30^ Proteins were labelled with all UniProt-identified substrates. Sequences with their donor substrates listed as a generic NDP-α-D-glucose donor substrate were excluded. Sequences with a donor substrate appearing less than three times in the dataset were also excluded, as at least three examples are needed for representation in the training, cross-validation, and test sets. This curation resulted in a dataset with seven unique donor substrates (GDP-α-d-mannose (GDP-Man), GDP-l-β-fucose (GDP-Fuc), UDP-α-d-galactose (UDP-Gal), UDP-α-d-glucose (UDP-Glc), UDP-α-d-glucuronic acid (UDP-Glcua), UDP-α-d-xylose (UDP-Xyl), and UDP-β-l-rhamnose (UDP-Rha)).

As the structural diversity of these enzymes is likely to cause inaccurate sequence alignments, only sections of the sequence corresponding to the characteristic Rossmann-like domains^9^ and catalytic regions were used in training. Each residue’s secondary structure assignment was made using AF2 structures for each sequence and PyMOL.^31^ While a Rossmann-like fold is generally considered to have six to seven β-strands in each sheet, many structures in the dataset contained less, therefore sheets of at least four strands were considered sufficient for our curation. Fourteen proteins from the dataset were excluded from this analysis for lack of clear Rossmann domains. This curation eventually resulted in 474 GT-B proteins (dataset hosted at https://github.com/samihennen/GTB_Substrate_Prediction/tree/main/Data) to be further split into training and test sets. The percent shared identity was calculated for all sequence pairs, with the test set sequences chosen if they contained less than 75% identity to any training sequences. This resulted in final training and test datasets of 381 and 93 sequences, respectively. The training dataset contained 14 GT-B families with seven distinct donors (SI Figure 2, Table 1). To further assess model robustness, additional test subsets were generated to exclude proteins of identity score cutoffs of 70%, 65%, 60%, 55%, and 50% with any training sequence.

### 2.2. Featurization

The reduced sequences were aligned with Clustal Omega to create a multiple sequence alignment (MSA).^32^ Highly gapped residue positions in the aligned sequences were removed from the MSA to select only the most relevant features, and those that might have a relationship to structure. The MSA used in previous ML work on GT-A fold enzymes was curated similarly, with positions of over 15% gaps instead removed.^14^ This cutoff was increased to 50% in this work, as the lower cutoff would result in few remaining positions in the highly dissimilar GT-B fold enzymes, resulting in 803 removed MSA positions. The MSA was then converted to a dataset of feature vectors representing each sequence, featurized for residue property values. Each residue within the MSA was featurized with AAIndex assigned values for hydrophobicity, residue volume, accessible surface area (ASA), polarity, and charge.^33^ Solvent accessible surface area (SASA) and categorical secondary structure values were also assigned for each residue from AF2 structures by BioPython and PyMOL, respectively.^31, 34^ ASA and SASA differ as SASA accounts for the position of the amino acid residue in the structure, while ASA is dependent only on residue identity. AF2 residues with confidence scores below 70% were assigned values of 0 for solvent accessible surface area and secondary structure. Feature values were normalized between 0 to 1 for each feature type to prevent overemphasis on features with larger values. SASA values were normalized relative to each structure. As most models only allow non-null feature values, gapped positions received values of 0. Finally, to remove redundant features, 496 features with high correlation (>90%) were removed. This refinement was performed as the unfiltered feature vectors could result in inefficient, overfit models. This curation resulted in 1905 features to be further refined during model training.

To maintain identical feature length between training and test sets, each test sequence was aligned to the training MSA. The test sequences and structures were then featurized for the same features used in the training set.

### 2.3. Model Training

Four classifier ML model types (Random Forest (RF), K-Nearest Neighbors (KNN), Gaussian Naïve Bayes (GNB), and Support Vector (SV)) from the Scikit-Learn Python package were trained for multi-label classification with the 381 sequences in the training set.^35^ An overview of the model featurization and training protocol is depicted in Figure 1.

**Figure 1.**
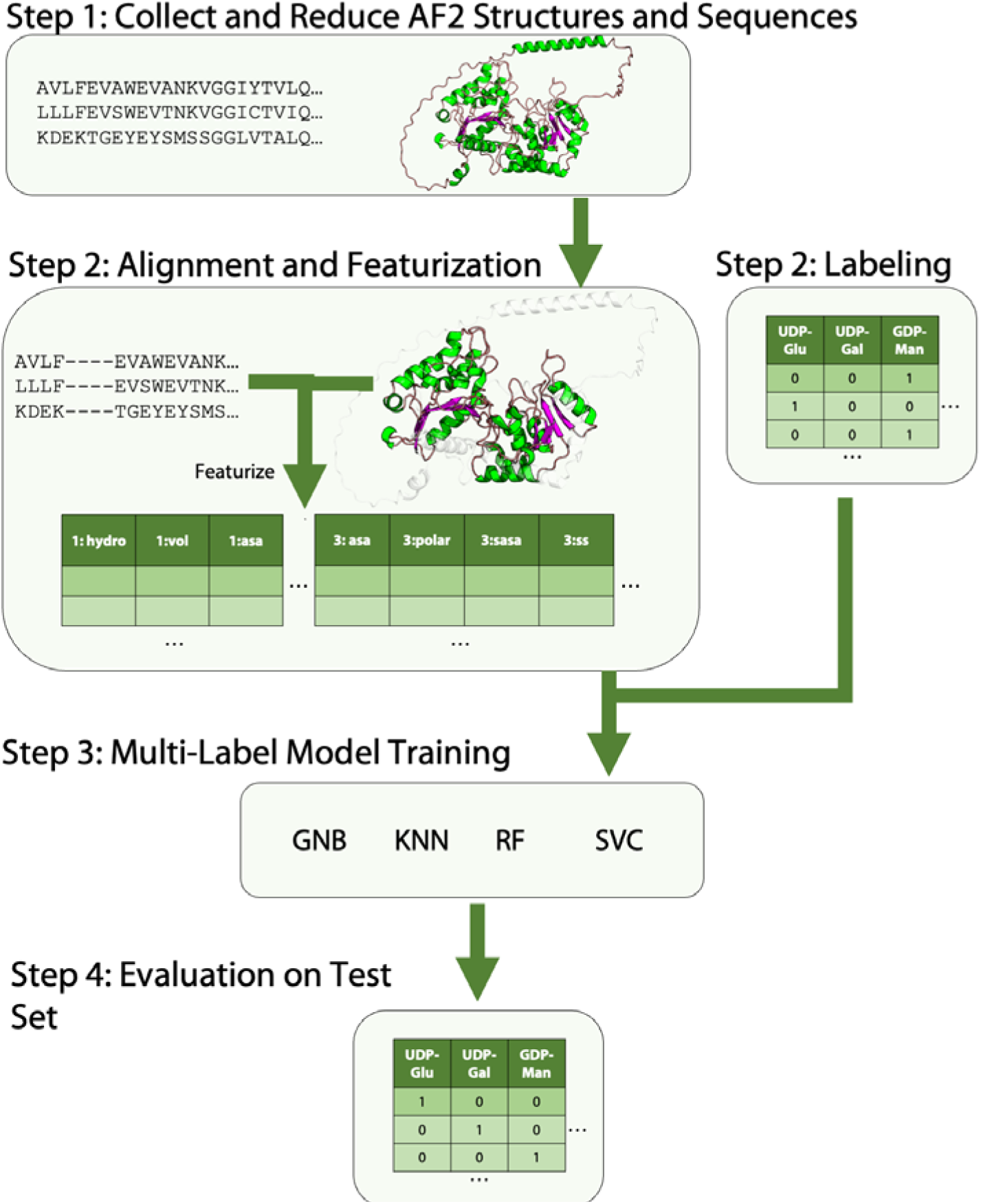
An overview of model featurization and training for GT-B fold glycosyltransferases. In Step 1, the AF2 structures were collected and reduced to the Rossmann-like domains. For Step 2, a multiple sequence alignment was constructed and sequences were featurized based on amino-acid properties such as hydrophobicity, volume, accessible surface area, polarity, charge, and structure-based amino acid properties - solvent accessible surface area and secondary structure. Highly gapped MSA positions and highly correlated positions in the featurized MSA were removed. The GTs were also labelled with the substrates to which they are confirmed to bind for prediction. Next, several model types were trained on feature subsets in Step 3, and their hyperparameters were tuned to identify optimal values. Finally, these models were assessed on the test set in Step 4.

Hyperparameters, such as the number of trees in an RF model, control a model’s complexity and can drastically alter performance, thus it is necessary to evaluate several combinations. Furthermore, different feature lengths should be evaluated to determine the minimum features needed for high accuracy to maximize efficiency and reduce overfitting. Therefore, hyperparameter tuning and feature selection was performed and optimized with leave-one-out cross-validation. Test set sequences were kept separate from those used for training and cross-validation for later performance assessment. F1 scores were averaged for each multi-label sample and used as the evaluation metric. To select the feature subset with optimal model performance, Scikit-Learn’s chi2 SelectKBest was used to select various feature subsets for model training.^36^ For each feature length between 50 and 1000, in multiples of 50, the model was trained with every combination of hyperparameters and assessed for performance. All hyperparameter search ranges can be found in SI Table 2.

### 2.4. Family Based Model

To ensure that the model was learning more significant relationships than simple family identifications, it was also trained with family number as its only feature and its cross-validation and test set performance compared to the model of higher complexity. The training and test sets were kept identical. All models (SVC, RF, KNN and GNB) were trained with this single-family feature and their hyperparameters tuned with an identical protocol as the more complex model.

### 2.5. Identifying and comparing conserved regions and binding sites

Residue positions used by the optimized model were assessed for consensus to elucidate relationships between conserved residues and substrate specificity. GT-B fold enzymes that bind UDP-Glc were considered here as it is a commonly observed donor substrate with known activity and is utilized by enzymes in several different families. As these families have dissimilar sequences (and structures), this evaluation was intended to explore the presence of structural components that might dictate substrate specificity. Residue positions whose features contributed to the optimized model were assessed for consensus of residue type (hydrophobic, polar, positive versus negatively charged) due to the minimal conservation between GT-B fold families. AutoDock Vina^37, 38^ blind-docking simulations were run on GT4, GT20 and GT28 candidate structures with UDP-Glc as the ligand. The GT structures were shortened to the Rossmann regions used in the ML models. 100 potential ligand poses were produced with the exhaustiveness parameter set to 320. High consensus residues were mapped onto the docked enzyme-substrate complexes to assess their role in binding.

### 2.6. Application to Other Genera

The trained and tuned models were used to predict substrate specificities of uncharacterized GT sequences from four dissimilar chlorophyte genera exclusive to the training dataset - *Populus, Spirodela, Chlamydomonas*, and *Eucalyptus.* While the training and testing sets where restricted to only Uniprot-reviewed sequences that are also classified as GT-Bs by CAZy, this curation resulted in a minimal number of uncharacterized sequences for the aforementioned genera. Therefore, we included Uniprot-unreviewed sequences and additional UniProt-classified GT-B fold enzymes in addition to the CAZy-classified GT-B sequences. Each protein sequence and structure was pared down to the Rossmann-like domains and the catalytic region. Any sequences without available AF2 structures, with AF2 structures of low confidence (residues < 70% confidence) or lacking clear Rossmann-like domains were excluded. The final sets for *Populus, Spirodela*, *Chlamydomonas,* and *Eucalyptus* included 308, 146, 162, and 375 enzymes respectively. After reducing the sequences and structures to the Rossmann-like domains, alignment to the training MSA and featurization, our optimized models were used to predict their potential donor substrate specificity.

## 3. Results

### 3.1. The characteristic Rossmann-like fold anchors plant GT-B structures

The GT-B fold was first described for the bacterial T4 β-glucosyltransferase^39^, and has since been found in many GT families including GT28, GT35 etc.^40^ The characteristic aspects of the GT-B fold consist of two Rossmann fold subdomains and a loop connecting them that plausibly acts as a hinge to mediate catalysis and specificity.^40^ The Rossman fold itself is known to be one of the most ancient, prevalent and functionally diverse protein folds that involve nucleoside-based cofactors^41, 42^ While experimental structures for GT-Bs from non-plant systems such as bacterial and animal kingdoms have been available for some time,^6, 43^ structural characterization of plant GT-Bs have been less forthcoming. Recently, the Rossmann-like fold was established as one of the hallmark features for the plant fucosyltransferase from *A. thaliana* classified as a GT37.^15^ With particular interest in plant-based GT-Bs and in anticipation of the challenges presented by the inherent structural diversity of GT-Bs, we analyzed the AF2 structure of all the sequences being considered in this study. We decided to parse the sequences to look for sections that correspond the characteristic Rossmann-like subdomains. Our analysis verified that almost all the sequences had characteristic Rossmann-like subdomains, with a few exceptions either too short or containing too few β-strands. This is illustrated in Figure 2 where these domains in candidate structures from 14 distinct GT-B families (of which nine are common families observed in plants) are highlighted. All of them have a N-and C-terminal Rossmann-like subdomains. The non-Rossmann-like fold domains of the structures are hidden for the sake of clarity. The GT-B binding and catalytic active site are likely situated at the interface of the two Rossmann-like subdomains.

**Figure 2.**
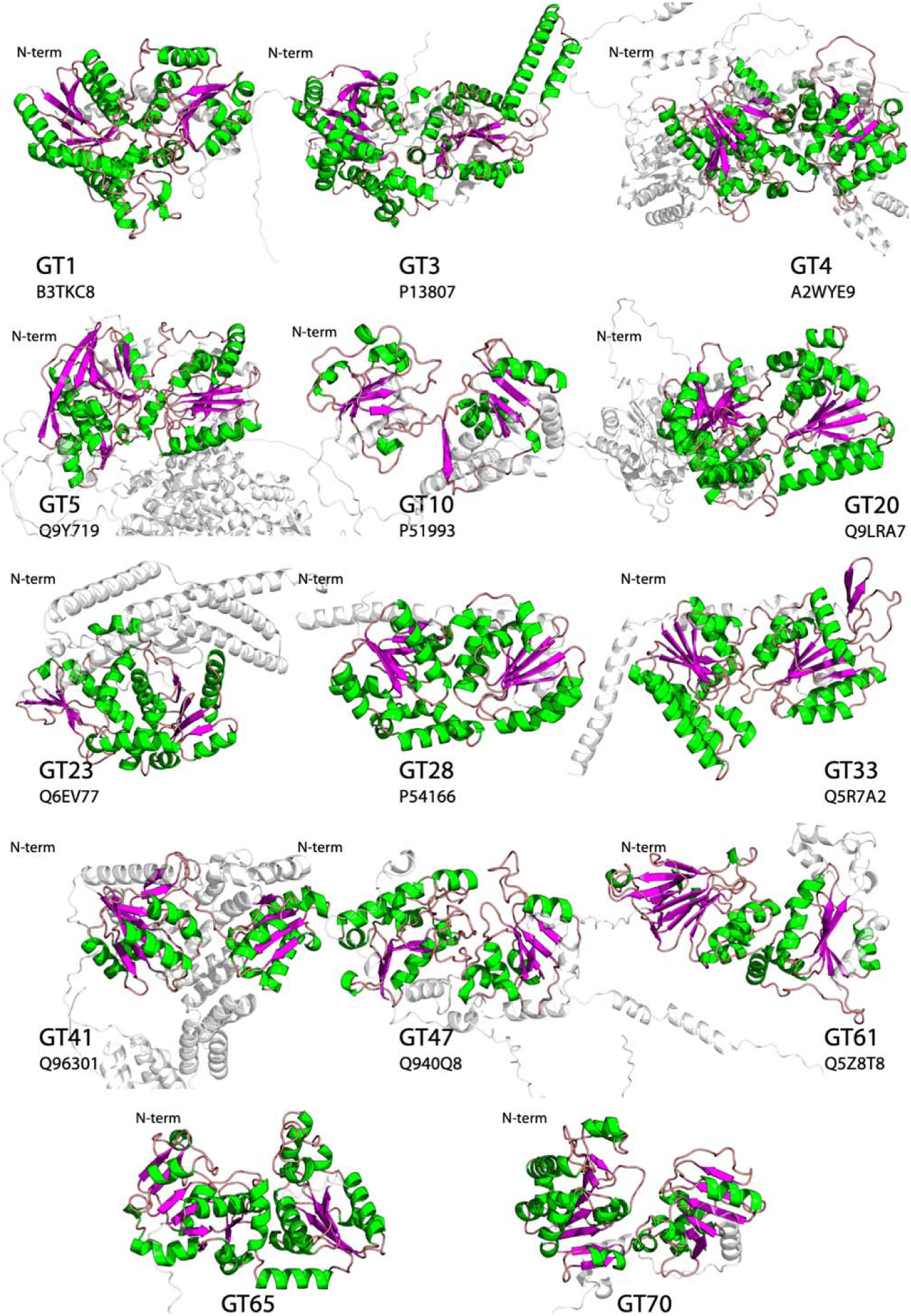
Representative structures for 14 plant GT-B families represented in the dataset (rendered using PyMOL), with β-sheets shown in magenta and α-helices in green. While there is significant diversity in the structure and size of these proteins, all structures contain the characteristic Rossman-like subdomains. The structures are all aligned such that the N-terminal subdomain is on the left while the C-terminal is on the right. Sections of each structure that are not part of the Rossman-like subdomains are depicted transparently for clarity.

### 3.2. Trained models yield high accuracy on cross-validation and test-sets

The ability of the four multi-label ML models to accurately predict the correct donor substrate specificity of the training set sequences was evaluated using cross-validation and test scores. All models achieved cross-validation F1 scores of at least 89%. As varying the feature lengths resulted in models with very similar cross-validation scores, the model with the best test score was chosen for further evaluation. The KNN model with 850 features had the best balance for cross-validation (95%) and test scores (85%) (Figure 3). As the dataset’s substrate representation is imbalanced, an additional metric - the Matthew’s Correlation Coefficient (MCC), was evaluated for each of the test set substrates to assess the model’s performance on all labels (Figure 3C). Finally, a confusion matrix was also generated for the test set predictions (Figure 3D). As this is a multi-label model, a false positive for a substrate was only considered if the true positive was not correctly predicted. The model achieved high accuracy for four of the seven substrates. The remaining three substrates, GDP-Fuc, UDP-Xyl, and UDP-Rha, with lower test set performance, show declining performance with fewer examples in the training set. The training set contained 42, 8, and 3 examples of these substrates, respectively. While GDP-Fuc had ample representation in the dataset, its lower MCC score of 69.4% may be owed to the diversity of the training set samples, representing four GT families. GDP-Man is similarly represented in the dataset, with 44 samples, from only two GT families and has a higher MCC score of 90%. The hyperparameters and feature lengths for the chosen models, with optimal cross-validation and test set scores, can be found in SI Table 3.

**Figure 3.**
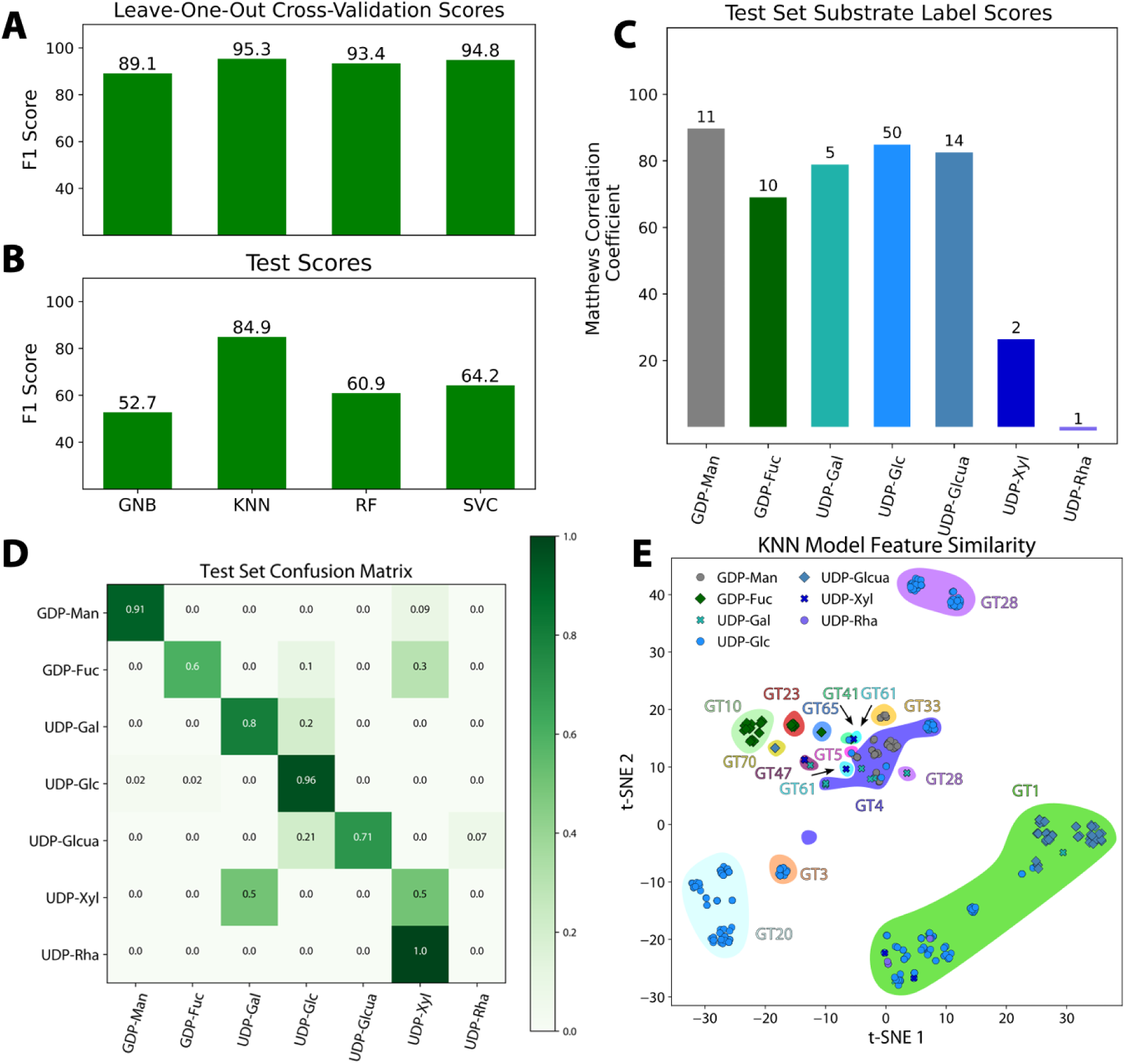
(A) F1 cross-validation scores and (B) test scores indicate that KNN model performs the best with the GNB model performing the worst. (C) The Matthews Correlation Coefficient scores reveal substrate-level accuracies for the best performing KNN model. The number of test set samples for each substrate is indicated. (d) A confusion matrix for the test set indicates substrates that are mis-classified by the model. (e) Dimensionality reduction of the 850 selected features from the KNN model with T-distributed Stochastic Neighbor Embedding (t-SNE) reveals unique family and substrate-based clustering.

### 3.3. Clustering reveals unique structural motifs within GT-B families

To visualize any trends in model classifications, a dimensionality reduction was conducted for the feature vectors from the KNN model using the T-distributed Stochastic Neighbor Embedding (t-SNE) method.^44^ In Figure 3E, each dot on the plot represents a substrate classification for a test sequence by the model, with shaded regions indicating the GT family to which the sequence belongs. There is significant clustering of GT sequences binding the same substrate, which might be central to the KNN model’s high accuracy. While much of the clustering is likely due to family identity, there are some distinctive clustering features. One of these is the fact that some sequences from the same family cluster in different locations based on substrate specificity as exemplified by GT28 sequences organizing into three clusters, two labelled with UDP-Glc and one with UDP-Gal. Notably, the UDP-Gal cluster is located nearer to the GT4 sequences that also have UDP-Gal specificity. The other interesting observation is that the GT1 sequences are a very dispersed cluster, likely due to great donor and acceptor diversity observed for these enzymes.^45^ An analysis of the acceptor substrates, curated in a similar fashion as the donor substrates (see Section 2.1), found GT1 enzymes that use UDP-Glc as a donor bind dozens of distinct acceptors, including several different benzoxazinoids and flavonols.

Notably, the t-SNE analysis shows several UDP-Glc sequences as clustering into distinct groups within their respective families, from GT1, GT20 and GT28. This clustering was further inspected by comparing structures from each cluster with significant structural differences found between the clusters (Figure 4). Two of the GT1 clusters (Figure 4A.2, 4A.3) are in close proximity in the t-SNE analysis (Figure 4A.1), and representative structures from each cluster show high similarity with minor differences in α helix locations. Conversely, a representative structure from a more distant cluster shows much more significant structural differences (Figure 4A.3). A similar assessment for the GT20 clusters shows similar structural differences, with two of the clusters containing additional β strands from the third cluster (Figure 4B.2-4B.4). The assessment for the GT28 clusters shows more subtle differences, most significantly in shorter β strands in one of the clusters (Figure 4C.2-4C.3). This analysis indicates that the 850 feature t-SNE analysis is able to encode structural differences even in the same GT families.

**Figure 4.**
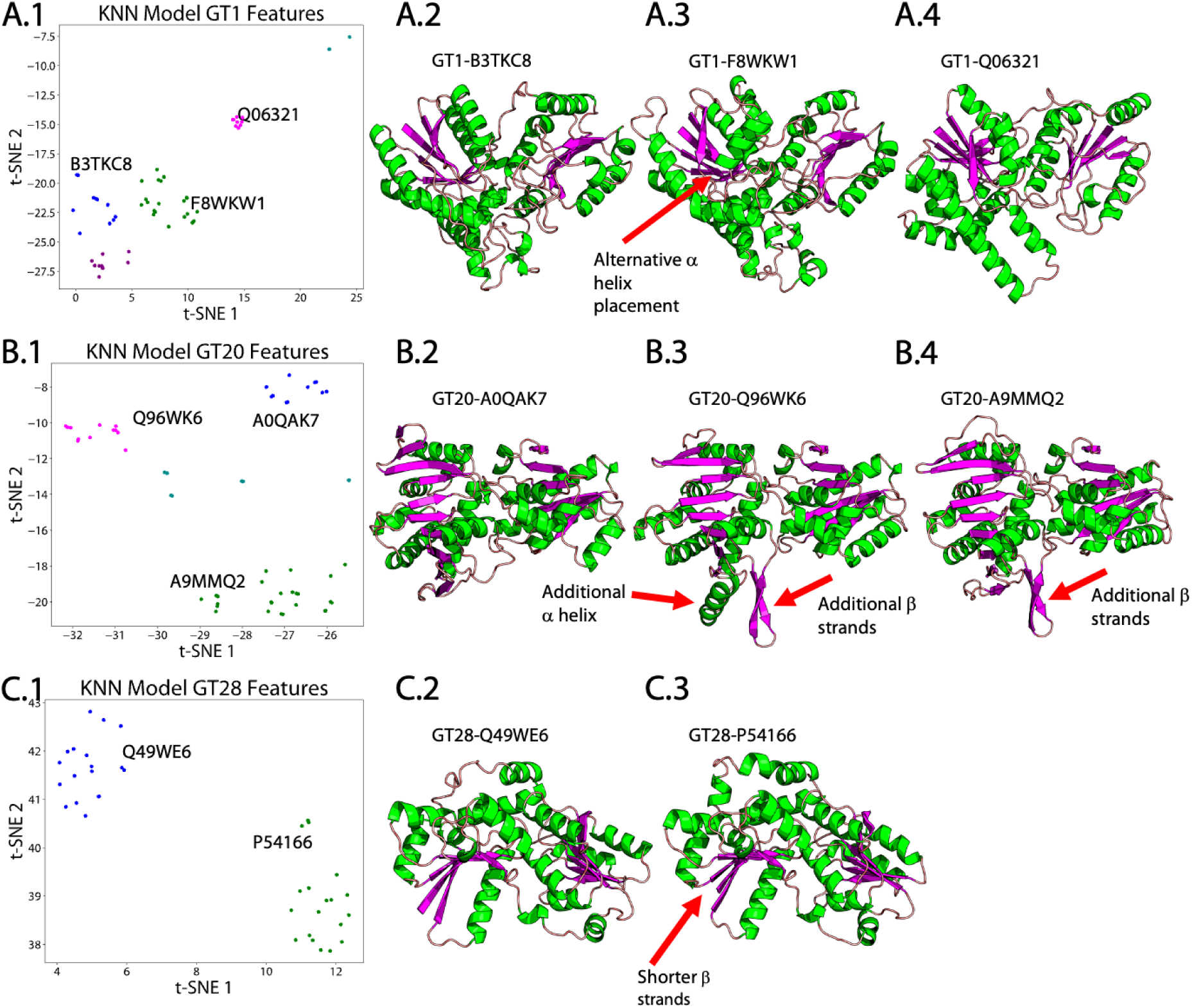
(A.1, B.1, C.1) t-SNE plots derived from Figure 3E focusing on specific regions is shown only for UDP-Glc binding sequences from families GT1, GT20, and GT28, respectively. (A.2-A.4, B.2-B.4, C.2-C.3) Structures from each cluster are shown with selected distinguishing structural features indicated.

To ensure that the accuracy of results are not driven by high sequence similarity between the testing and training datasets, we created testing data subsets that excluded proteins that share high sequence identity with the training dataset sequences and evaluated the KNN model on these subsets. Unsurprisingly, the model accuracy declines on subsets with lower shared identity. MCC scores were again calculated for each substrate in the given set and the average shown in Table 1. The overall F1 score declines at lower shared identity values but remains ∼76% at only 50% shared identity suggesting that the model accuracies not due to high-sequence similarities in the training set. The average MCC score similarly declines at lower identity cutoffs, with the lower performance largely due to a loss of accuracy in GDP-Fuc predictions.

**Table 1:**
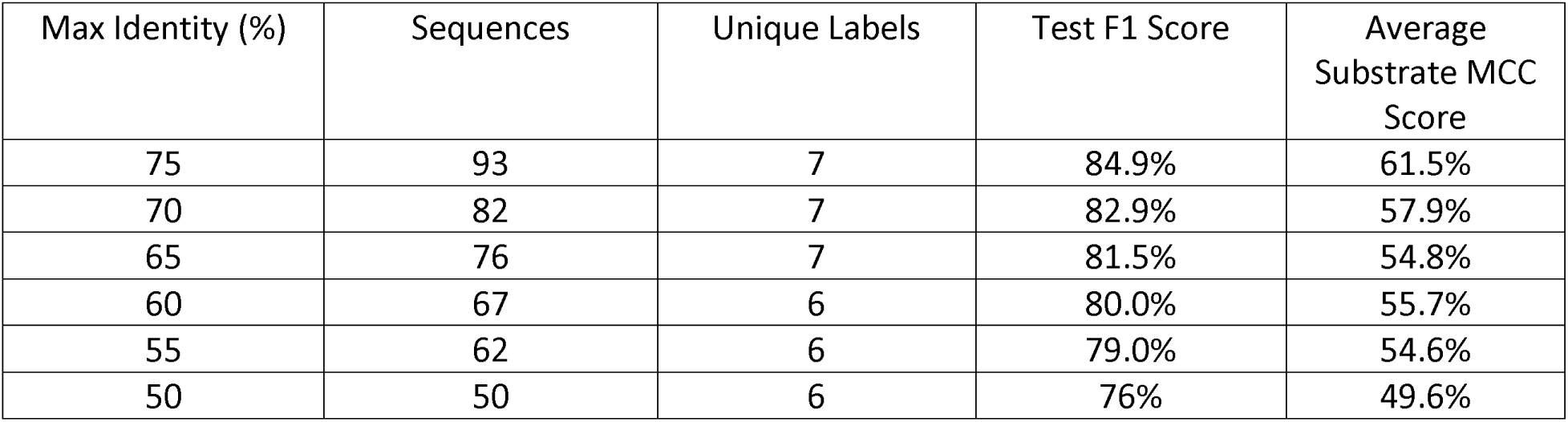
The KNN donor prediction F1 scores on the test set subsets are shown, where proteins sequences above the full sequence identity similarity cutoffs to any training set sequences were excluded.

Furthermore, to ensure that our 850-feature KNN model’s substrate classifications are not merely family-based, we also built single-feature models and trained them with just the family identifier as its feature (SI Figure 3). This analysis verified the importance of including the complete feature set in model training, with the best test set scores of 84.9% for the complete feature model and only 62.4% for the family-based model.

### 3.4. Relating conserved residues to structure

To assess whether the features used by the ML model could elucidate common structural features between dissimilar families, residues involved in features used by the KNN model were evaluated for consensus. Enzymes active on UDP-Glc were chosen for this evaluation, as these enzymes comprise 204 of the 381 total sequences and represent seven distinct families. GT-B fold families GT1, GT3, GT4, GT5, GT20, GT28, and GT41 participate in UDP-Glc binding and are represented by 72, 14, 16, 1, 62, 37 and 2 sequences respectively. Conservation amongst residues contributing to the 850 substrate-specificity determining features of the KNN model was assessed with a 90% cutoff within each family as well as amongst all families. While no residues had >90% conservation amongst all seven families, there was some conservation amongst the three families-GT4, GT20 and GT28, with 10 residues conserved at over 90% (Table 2). Figure 5A depicts a Venn diagram that shows the number of conserved residues within each family, between any two families and amongst all three families. For example, within 16 GT4 family enzymes that bind UDP-Glc, there are 85 positions conserved in the 850 features used by the KNN model. A similar analysis of 62 GT20 sequences that bind UDP-Glc reveals 81 conserved positions that belong to the KNN feature set. Similarly, amongst the 37 GT28 sequences, there are 133 conserved MSA positions.

**Table 2:** The residues corresponding to the MSA positions with property consensus over 90% within sequences from GT families 4, 20 and 28, and is restricted to enzymes that utilize UDP-Glc as a donor. NC indicates that no conservation was observed in that particular structure, while other structures from the same family do have conservation at that MSA position.

|  | MSA Position |  |  |  |  |  |  |  |  |  |
| --- | --- | --- | --- | --- | --- | --- | --- | --- | --- | --- |
|  | 676 | 683 | 806 | 837 | 913 | 1012 | 1015 | 1041 | 1068 | 1113 |
| A2WYE9<br>(GT4) | V453 | F460 | L508 | I517 | I539 | I587 | L590 | L609 | A615 | L640 |
| Q9LRA7<br>(GT20) | I306 | L311 | V340 | I349 | I375 | NC | Y427 | L445 | I451 | V477 |
| P54166<br>(GT28) | I179 | F183 | A209 | V218 | V236 | I267 | L270 | I284 | NC | NC |

**Figure 5.**
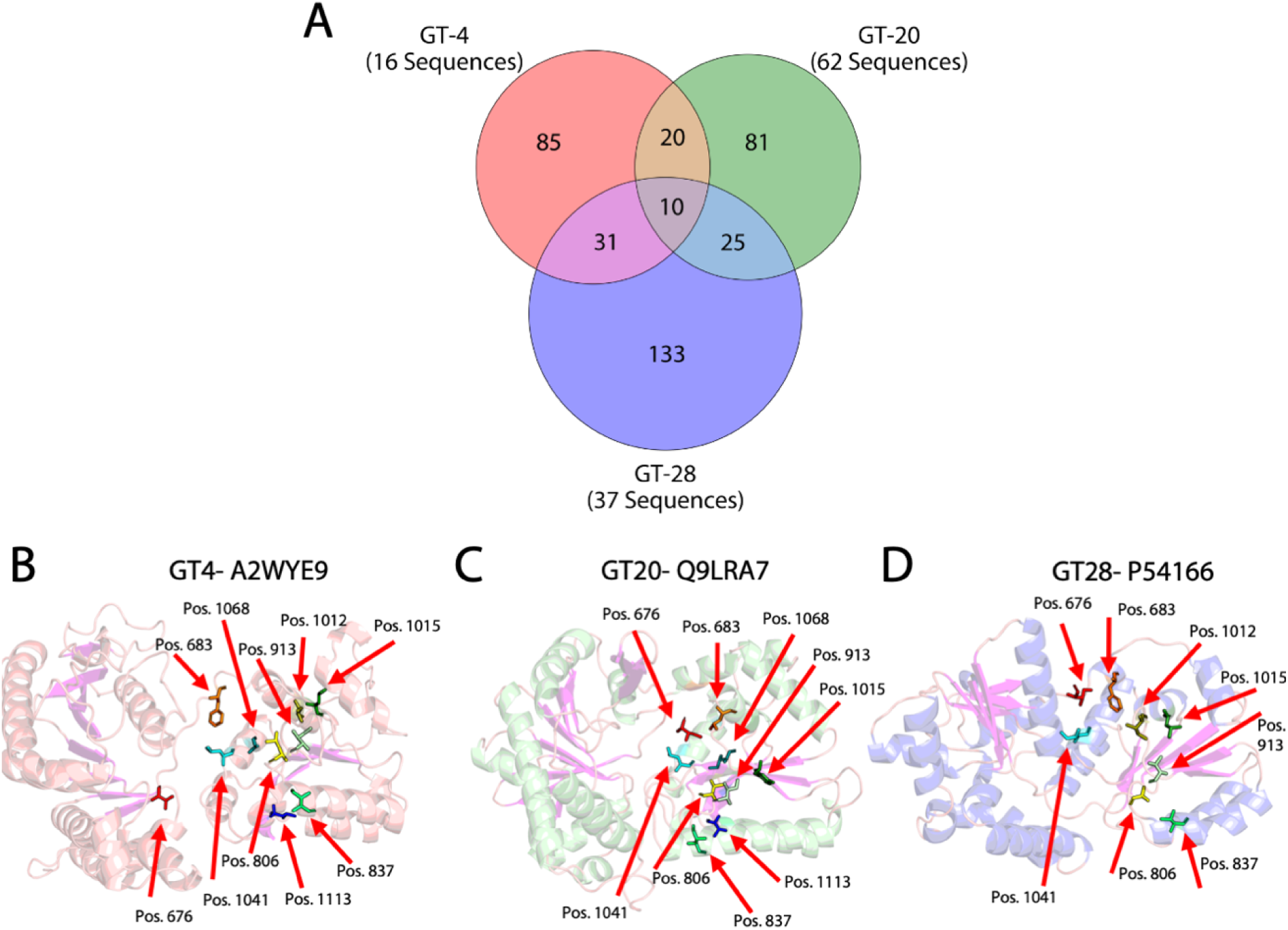
(A) Venn diagram showing the distribution of residues (MSA positions) that are part of the 850 features used by the KNN model and are involved in detecting substrate specificities for GT4, GT20 and GT28 families constituting 16, 62 and 37 sequences respectively. The overlapped regions indicate the portion of residues (MSA positions) that are commonly conserved amongst any two and all three families for the same property. (B-D) The α-helices are color-coded based on the Venn diagram with the GT4 structures depicted in salmon, GT20 in pale green and GT28 structures in blue. (B-D) Structural regions used in model training are show with the 10 conserved MSA positions common to all three families depicted in licorice representations. Ten, nine, and eight of these positions are conserved in the example GT4, GT20 and GT28 structures respectively.

Since the GT4, GT20 and GT28 families share UDP-Glc as a common nucleotide sugar donor substrate and exhibit structural similarity in their Rossmann domains, it is not far-fetched to imagine that the 10 residue positions within the KNN features with consensus above 90% (all for hydrophobicity) in each family might reveal common structural motifs responsible for the selectivity for this substrate (Figure 5A). Residues corresponding to these positions are listed in Table 2 for a representative structure from each family. Ten, nine and eight of these positions show consensus in the chosen representative structures for families GT4, GT20 and GT28, respectively. These residues exhibit similar placement in all these structures, with the majority located near the C-terminal Rossmann region (Figure 5B, 5C, 5D).

The direct involvement of these conserved residues in substrate binding was investigated by docking UDP-Glc to each of these enzymes using Autodock Vina. As expected, all docked poses (shown in SI Figure 4) for the three candidate enzymes were observed to be at the interface between the N-terminal and C-terminal Rossmann-like subdomains, which contain the presumed active site. While five positions in the GT28 structure are observed to be at the putative substrate binding site, only one position in the GT20 and no positions for GT4 structure are observed. Therefore, a significant relationship between the conserved residues and their role in substrate binding could not be established for this case.

### 3.5. Model extension to uncharacterized sequences from other plant genera

Finally, the optimized KNN model was applied to uncharacterized GT-B fold enzyme sequences from distinct plant genera to predict their substrate specificities, including *Populus, Spirodela*, *Chlamydomonas,* and *Eucalyptus* (Figure 6). The datasets contain 308, 146, 162, and 375 enzyme sequences, respectively. These genera were chosen in part due to their distinct physiology, a consequence of different carbohydrate composition profiles of their cell walls. The species-specific distribution of families for this dataset is listed in SI Table 4. While the KNN model can generate predictions for multiple substrates, no enzymes in any of the sets were predicted to be substrate promiscuous. The predictions for each genera potentially reflects a different distribution of substrates, with UDP-Glc being one of the most common donors predicted for all genera except *Chlamydomonas* GT-Bs. Furthermore, the most abundant donors predicted for *Chlamydomonas* GT-Bs are GDP-Fuc and UDP-Xyl, potentially indicating distinct carbohydrate profiles that are species-specific. It must be noted a limitation of this analysis is that the model can only predict the seven donor nucleotide-sugars that it has been trained on. It is likely that the some of the sequences that have been considered here might indeed be active on other donors that haven’t been considered in this study.

**Figure 6.**
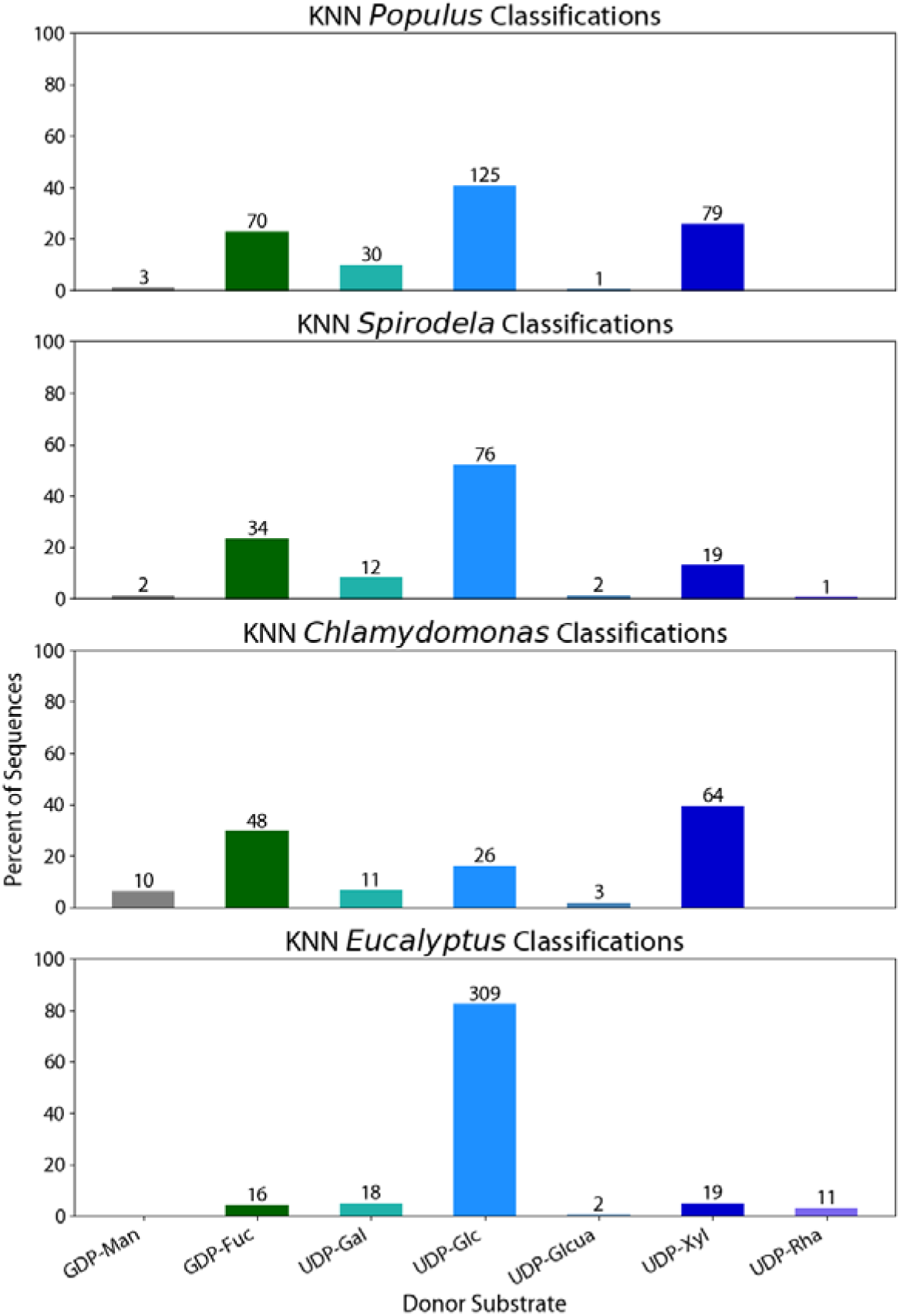
Application of the optimized KNN model to uncharacterized GT-B fold enzymes in diverse plants. Donor substrate classifications for the uncharacterized proteins from *Populus, Spirodela, Chlamydomona* and *Eucalyptus* are shown as the fraction of total sequences in the species dataset.

## 4. Discussion

Understanding the substrate scope and specificity of glycosyl transferases is crucial for our understanding of natural biosynthetic mechanisms of carbohydrate polymer synthesis. In this work, we develop and evaluate the efficacy of ML classifier models for the prediction of nucleotide donor substrates of uncharacterized GT-B fold enzymes. A major challenge we encountered was the curation of existing experimental data – namely the annotation of sequences and their activity for relevant nucleotide-sugar donor substrates. We collated an extensive dataset of GT-B sequences with experimentally characterized activities on seven donor nucleotide sugars. This data was used to train four different classifier models, amongst which the KNN classifier demonstrated the best performance for predictions on the training set, while also being generalizable to the test set. The feature vectors used by the KNN model formed the basis for further dimensionality reduction and residue conservation analyses. The dimensionality reduction analysis revealed both substrate-based and family-based clustering of sequences. We also noticed multiple sub-clusters for sequences from the same family (GT1, GT20 and GT28) and with activity on the same substrate (UDP-Glc). Structural analysis of sequences within these sub-clusters revealed subtle differences and indicated unique structural motifs for these subclusters. Furthermore, we assessed the KNN model’s feature-set for its ability to relate protein sequence to conserved structural regions and binding sites. While some limited structural consensus was revealed, our analysis did not reveal a strong relationship between the suggested binding sites and regions of high consensus.

Previous work has successfully used similar substrate activity classifiers, individually or as ensembles, for a variety of enzyme classes like bacterial nitrilases, thiolases and the GT1 family which adopts the GT-B fold.^23, 46, 47^ A general enzyme prediction model has also been developed using neural networks. However, its accuracy notably declines on substrates not well represented in its training set, such as the donor substrates common to GT-Bs, demonstrating the need for more specific enzyme class models.^48^ Thus, RF, SVC and KNN models were applied to our dataset of 381 samples, with an additional challenge presented by these samples being derived from highly dissimilar families adopting the GT-B fold. While enzymes from these families share a distinct structural Rossmann fold motif, they exhibit high sequence dissimilarity. Therefore, we incorporated structural data as well, through both reducing sequences to only the Rossmann domains and featurizing AlphaFold2 structures for solvent secondary surface area and secondary structure. Notably, an additional finding of this work is the inclusion of SASA and secondary structure features made little difference in test accuracy (SI Fig 5). Liu et al made a similar discovery in their work on residue pKa prediction, where they also found SASA data to make little difference in accuracy.^49^ A potential reason for this lack of improvement may be that the AlphaFold2 structural data is a function of sequence and is already incorporated into the feature data. Additional features instead derived from molecular dynamics simulations and docking simulations, such as RMSF and binding affinity data, may provide necessary physics-based information for substrate classification.^47, 50^

A phylogenetic analysis reveals one of the inherent complexities involved in resolving substrate-specificity determining factors in GT-Bs. Figure 7A shows the phylogenetic tree as generated from the full-length sequence alignment of all 381 training sequences using the NJ method.^51^ There is a clear family-based ordering of the sequences in this phylogenetic tree. However, an overlay of the substrate specificities on the same tree (Figure 7B) reveals that there are families that act on multiple substrates (eg: GT1), and there are multiple families that act on the same substrate (eg. UDP-Glc being acted on by GT1s, GT4s and GT20s). This clearly is consistent with experimental data that substrate-specificity is not family-based among GTs. Other possible reasons for lack of a relationship between conserved residues and binding sites include allosteric effects from these regions, or simply that common features among these proteins are not necessarily related to binding. Alternative ML model frameworks which more directly incorporate structural information, such as graph neural networks, may have more success in correlating the identified important residues with the substrate binding site.^52^

**Figure 7.**
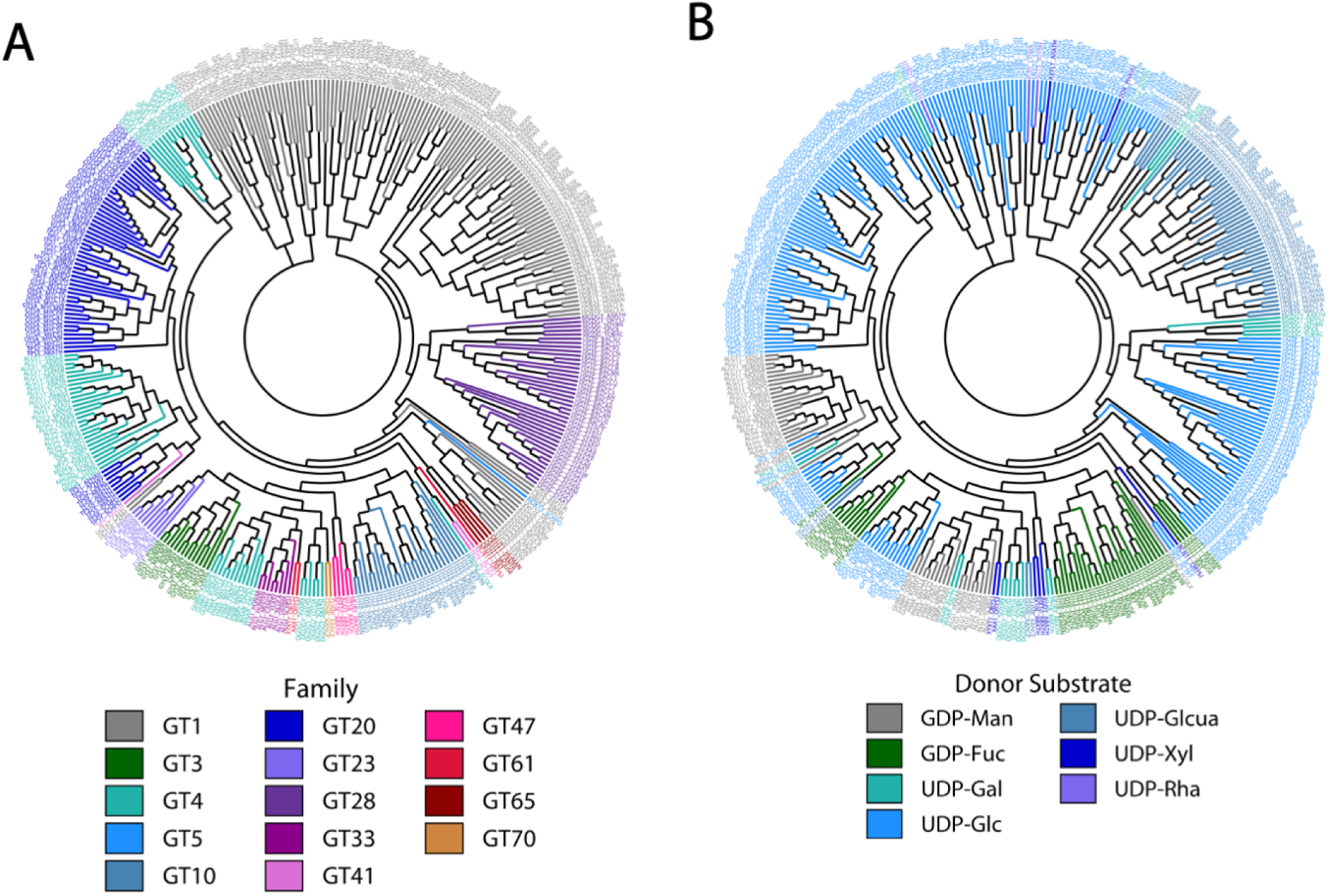
A phylogenetic tree constructed from the 381 full-length training sequences. (a) shows this tree overlayed with a single donor substrate for each sequence. (b) shows this tree overlayed with the GT family identification for each sequence.

A further limitation of this work concerns the training data quality. Many of the training enzymes may in fact have activity on several additional donor substrates, but this has not been experimentally studied, and therefore cannot be included in the substrate labels. The enzymes in this dataset had only one substrate listed on the database, but many may very well be less selective polyspecific enzymes. Despite this limitation, most enzymes are listed with the substrate used for their primary function(s), and so the KNN model would remain accurate in predicting the substrates for which the enzyme has the most activity.

While the models trained in this work focus exclusively on donor substrate prediction, extending these models for acceptor prediction is an important consideration for future work. However, this portends some considerable challenges including, due to inconsistent acceptor substrate labelling on databases and the ability of GTs to act on acceptor substrates with varying architectures.^53^

## 5. Conclusion

GT-B substrate specificity has been a challenge to characterize for diverse families due to a lack of shared sequence similarity, limited curated substrate specificity data and a paucity of structural information. However, with increased availability of sequences and predicted protein structures, ML tools that can reliably produce accurate predictions for the donor substrates of these proteins must be developed. Our study demonstrates that (1) current classifier ML models may be adapted to include structural data on these enzymes and (2) can predict substrate specificities for GT-Bs with reasonable accuracies but (3) interpretability does not allow for direct elucidation of structural features and (4) they are still severely limited by the paucity of curated biochemical and structural data on these enzymes. In the future, we envisage the development of novel ML architectures for these predictive models that will be adapted according to the nature of available characterized experimental and physics-based modelling data on these enzymes.

## Acknowledgments

This work was supported in part by the U.S. Department of Energy, Office of Science, Office of Workforce Development for Teachers and Scientists (WDTS) under the Science Undergraduate Laboratory Internship (SULI) program. This work was authored in part by the Alliance for Sustainable Energy, LLC, the manager and operator of the National Renewable Energy Laboratory for the U.S. Department of Energy (DOE) under contract no. DE-AC36-08GO28308. This research was supported by the U.S. Department of Energy, Office of Science, Biological and Environmental Research, Genomic Science Program grant no. DE-SC0023223. We are also grateful for the access and use of NREL’s Computational Sciences’ resources (Eagle) supported by the DOE Office of EERE under contract no. DE-AC36-08GO28308.

## Author Contributions

VB conceptualized the project, assisted in model development and data-analysis and visualization. SH conceptualized the project, curated the data and developed the ML models, performed data-analysis and visualization. VB, SH, YB and BU wrote the manuscript.

## Data and Software Availability

The code and datasets are available at: https://github.com/samihennen/GTB_Substrate_Prediction.git

