## Supplementary Information for "Decoding substrate specificity determining factors in glycosyltransferase-B enzymes – Insights from machine learning models"

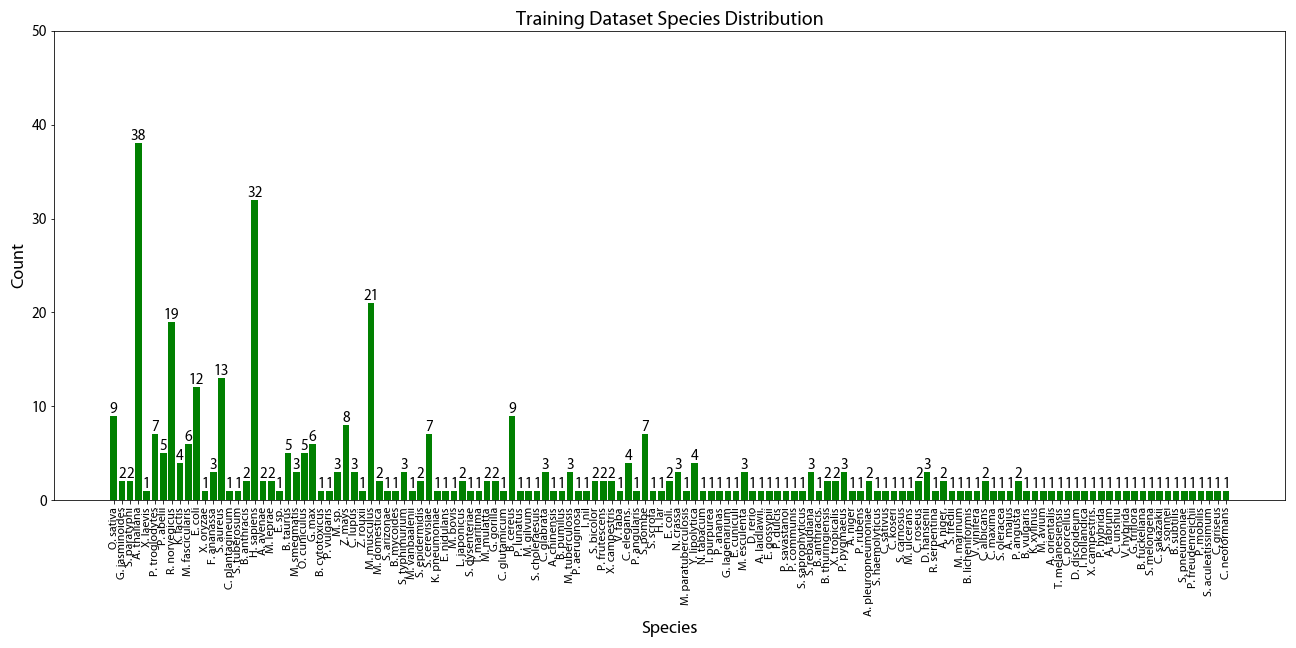

**SI Figure 1.** Distribution of sequences in the training dataset amongst plant and animal species.

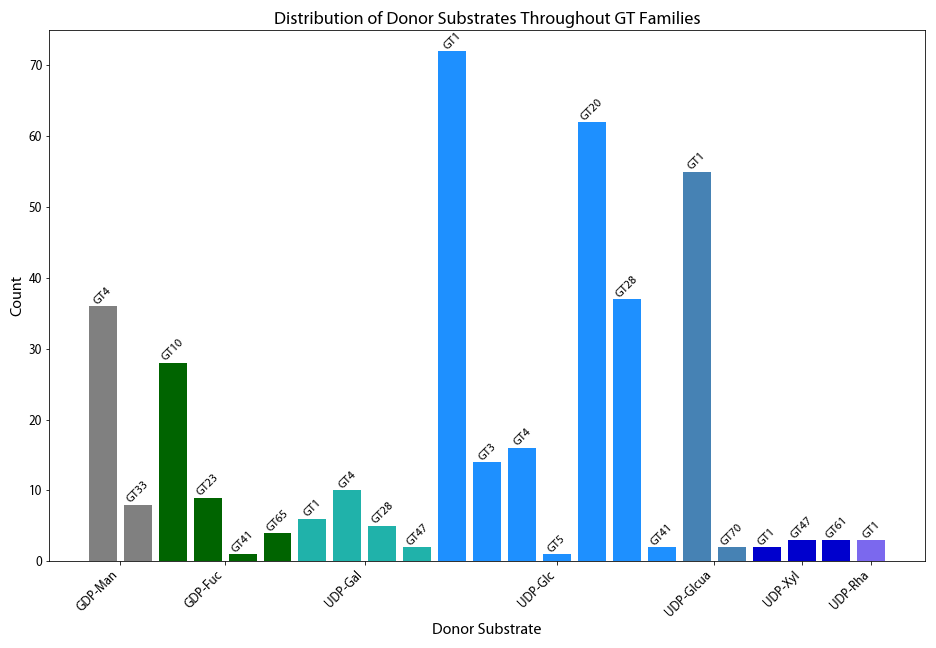

**SI Figure 2.** Number of sequences (and their family identity) in the training and test datasets with known activity for each nucleotide sugar donor substrate. Each bar denotes the number of sequences from a family with the color denoting its nucleotide-sugar activity.

**SI Table 1:** The number of sequences per family in the training dataset is shown below.

| **Sequences** | **Family** | **Donor Substrates** |
| --- | --- | --- |
| 138 | GT1 | UDP-α-D-galactose, UDP-α-D-glucose, UDP-α-D-glucuronate, UDP-α-D-xylose, UDP-β-L-rhamnose |
| 62 | GT20 | UDP-α-D-glucose |
| 62 | GT4 | GDP-α-D-mannose, UDP-α-D-galactose, UDP-α-D-glucose |
| 42 | GT28 | UDP-α-D-galactose, UDP-α-D-glucose |
| 28 | GT10 | GDP-β-L-fucose |
| 14 | GT3 | UDP-α-D-glucose |
| 9 | GT23 | GDP-β-L-fucose |
| 8 | GT33 | GDP-α-D-mannose |
| 5 | GT47 | UDP-α-D-galactose, UDP-α-D-xylose |
| 4 | GT65 | GDP-β-L-fucose |
| 3 | GT61 | UDP-α-D-xylose |
| 3 | GT41 | GDP-β-L-fucose, UDP-α-D-glucose |
| 2 | GT70 | UDP-α-D-glucuronate |
| 1 | GT5 | UDP-α-D-glucose |

**SI Table 2:** All model hyperparameters used in the training grid search.

| **Model Type** | **Hyperparameter Ranges** | | | |
| --- | --- | --- | --- | --- |
| **Gaussian Naïve Bayes** | *Variance Smoothing* |  |  |  |
|  | 100 points in logspace (0,-9) |  |  |  |
| **K-Nearest Neighbors** | *Number of Neighbors* | *Power Parameter* | *Weights* |  |
|  | 1-10 | 1, 2 | Uniform, Distance |  |
| **Random Forest** | *Number of Trees* | *Maximum Features per Split* | *Criterion* | *Class Weight* |
|  | 20-200, in multiples of 20 | 0.1 ,0.3, 0.5 | Gini, Entropy | Balanced, Balanced Subsample |
| **Support Vector** | *Regularization Parameter* | *Maximum Iterations* | *Kernel* | *Gamma* |
|  | 0.1, 1, 10 | 10, 50, 100 | Radial Basis Function, Linear, Polynomial | 1, 0.1, 0.01 |

**SI Table 3:** Optimal hyperparameters and feature lengths of all models found in the training grid search.

| **Model Type** | **Hyperparameter Ranges** | | | |  |
| --- | --- | --- | --- | --- | --- |
| **Gaussian Naïve Bayes** | *Variance Smoothing* |  |  |  | *Feature Length* |
|  | 0.0187 |  |  |  | 800 |
| **K-Nearest Neighbors** | *Number of Neighbors* | *Power Parameter* | *Weights* |  | *Feature Length* |
|  | 1 | 1 | Uniform |  | 850 |
| **Random Forest** | *Number of Trees* | *Maximum Features per Split* | *Criterion* | *Class Weight* | *Feature Length* |
|  | 60 | 0.5 | Gini | Balanced | 450 |
| **Support Vector** | *Regularization Parameter* | *Maximum Iterations* | *Kernel* | *Gamma* | *Feature Length* |
|  | 0.1 | 50 | Linear | 1 | 400 |

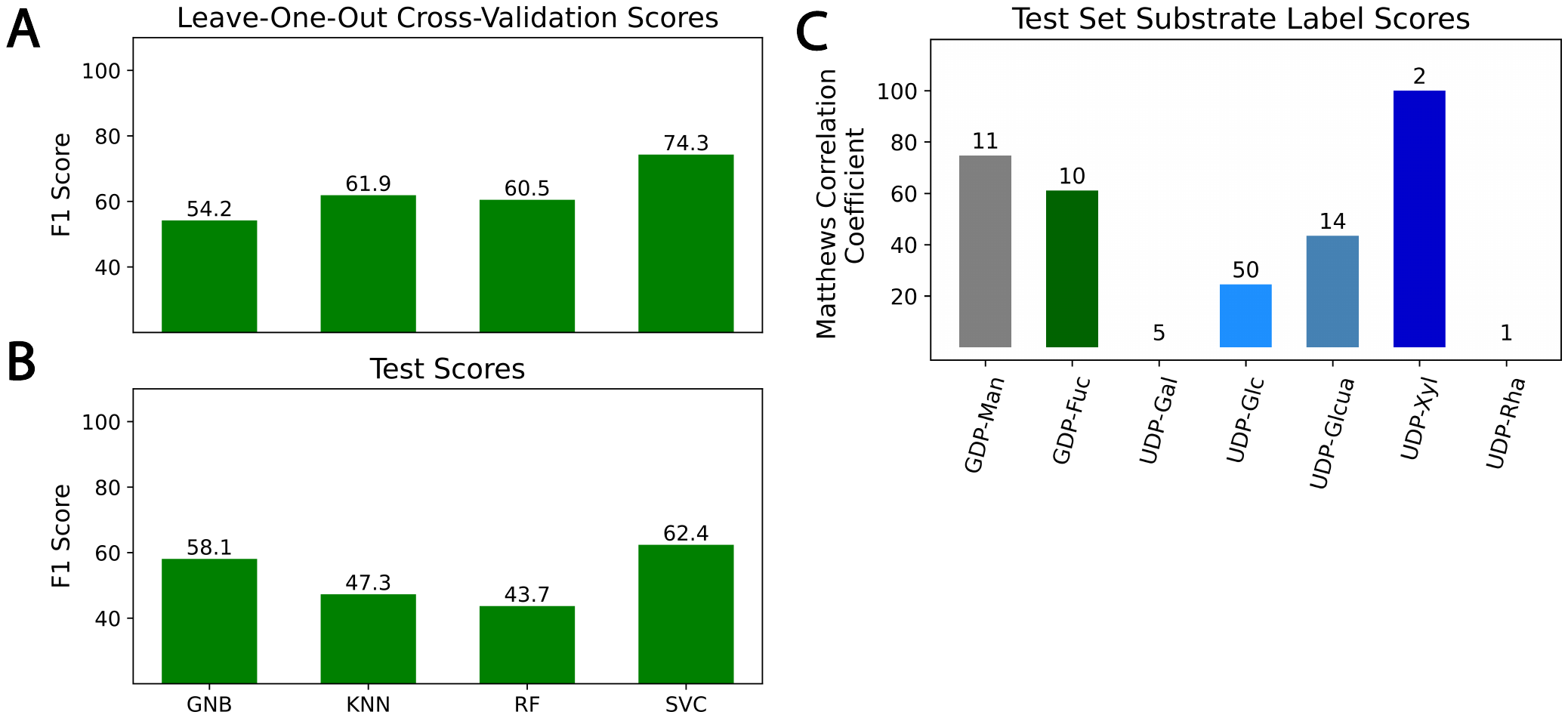

**SI Figure 3.** F1 cross-validation (a) and test scores (b) from all models trained on only the family number are shown. (c) The Matthews Correlation Coefficient scores of the best performing SVC model for each test set substrate are also shown. As expected, the cross-validation scores of all family-based models are lower than their counterpart models generated with additional features. The best test set score of 62.4% for the SVC model, is lower than the best test set score of 84.9% for the KNN model (Figure 3B) built with the complete feature set (Figure 3).

The individual substrate MCC scores are also lower than in the more complex model, showing similar or lower scores on all substrates except UDP-β-l-xylose. The high performance on this single substrate is notable, as it has higher accuracy than in the more complex model. Nonetheless, the poor performance on the additional substrates shows this family-based model’s inability to predict nucleotide sugar donor substrates.

**SI Figure 4.** (A-C) UDP-α-d-glucose was docked to the truncated representative structures A2WYE7, Q9LRA7, and P54166 from families GT4, GT20, and GT28, respectively. The top ranked pose for each structure is shown, along with residues found to be highly conserved (by residue type) within these families.

**SI Table 4:** The family distribution of uncharacterized sequences from distinct plant genera datasets.

|  | **Genera** | | | |
| --- | --- | --- | --- | --- |
| **Family** | ***Populus*** | ***Spirodela*** | ***Eucalyptus*** | ***Chlamydomonas*** |
| GT1 | 91 | 73 | 3 | 324 |
| GT4 | 0 | 2 | 4 | 0 |
| GT5 | 1 | 0 | 3 | 0 |
| GT10 | 0 | 0 | 1 | 0 |
| GT28 | 13 | 2 | 7 | 7 |
| GT37 | 28 | 13 | 1 | 3 |
| GT41 | 12 | 3 | 1 | 2 |
| GT47 | 147 | 49 | 139 | 35 |
| GT61 | 2 | 0 | 0 | 0 |
| GT92 | 14 | 4 | 3 | 4 |

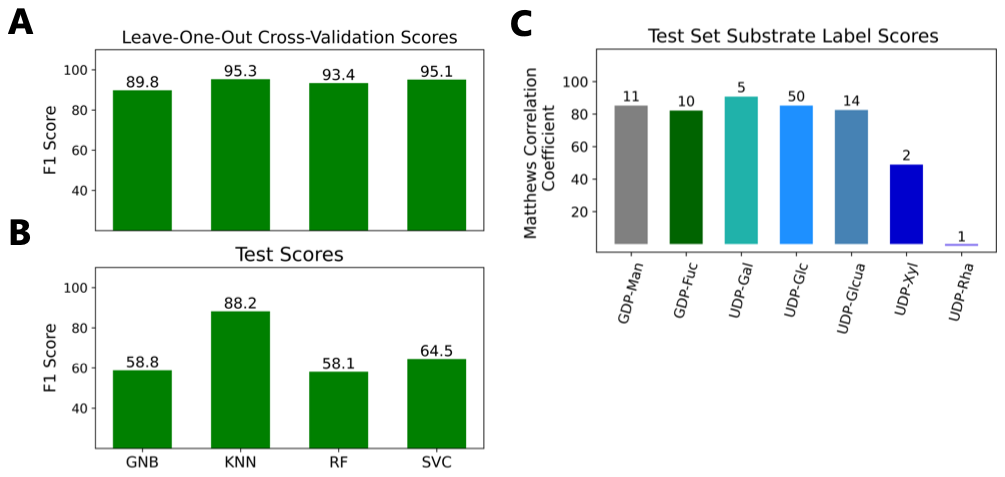

**SI Figure 5.** F1 cross-validation (a) and test scores (b) from all models trained without solvent accessible surface area and secondar structure values are shown. (c) The Matthews Correlation Coefficient scores of the best performing KNN model for each test set substrate are also shown.
